# A trade-off between error and synchrony when using temporal codes

**DOI:** 10.1101/309427

**Authors:** Erik J Peterson, Bradley Voytek

**Affiliations:** Department of Cognitive Science; University of California, San Diego, 9500 Gilman Drive La Jolla, CA, 92093; Neurosciences Graduate Program; University of California, San Diego, 9500 Gilman Drive La Jolla, CA, 92093; Halıcıoğlu Data Science Institute, University of California, San Diego, 9500 Gilman Drive La Jolla, CA, 92093; Pasteur Labs ISI

**Author notes:** EJP and BV designed the study and wrote manuscript. EJP implemented the model and conducted the analysis.

## Abstract

Neural oscillations can improve the fidelity of neural coding by grouping action potentials into synchronous windows of activity but this same effect can interfere with coding when action potentials become “over-synchronized”. Diseases ranging from Parkinson’s to epilepsy suggest such over-synchronization can lead to pathological outcomes, but the precise boundary separating healthy from pathological synchrony remains an open theoretical problem. In this paper, we focus on measuring the costs of translating from an aperiodic code to a rhythmic one and use the errors introduced in this translation to predict the rise of pathological results. We study a simple model of entrainment featuring a pacemaker population coupled to biophysical neurons. This model shows that “error” in individual cells’ computations can be traded for population-level synchronization of spike-times. But in this model error and synchronization are not traded linearly, but nonlinearly. The bulk of synchronization happens early with relatively low error. To predict this phenomenon we conceive of “voltage budget analysis”, where small time windows of membrane voltage in single cells can be partitioned into “oscillatory” and “computational”‘ terms. By comparing these terms we discover a set of inequalities that align with an inflection point in the curve of measured errors. In particular, when the entrainment and computational voltage terms are equal, the error curve plateaus. We show this point serves as a reliable natural boundary to define pathological synchrony in neurons. We also derive optimal algorithms for exchanging computational error with population synchrony.

*New and Noteworthy*. We establish exact conditions for when rhythmic entrainment of precise spike-times in a neural population will improve or harm it’s ability to communicate.

---

Rhythmic entrainment is a common feature of many biological systems, but complete synchronization in these same systems is often undesirable and even pathological to their function (1). This can be conceptually illustrated by neural oscillations, where a totally unsynchronized neuronal population might lack communication capacity whereas a perfectly synchronized population might lack computation capacity (2). The biological reality lies in between, where moderate oscillations coordinate the firing of many individual neurons, creating synchronous windows, and therefore enhancing the population’s communication (3). Temporally grouping action potentials in this manner improves signal to noise (4) and increases the number of coincident firing events (5, 6), driving learning at individual synapses (7–9). In many circuits though complete synchronization leads to pathological outcomes as in Parkinson’s disease (10) and in epilepsy (11, 12). A question then is, what is the optimal level of synchrony for a given neural circuit?

To begin to understand this problem more completely here, in this paper, we imagine a population where each neuron receives the same spiking input and responds to this input according to the incoming synaptic weights, and it’s own membrane dynamics. We eliminate recurrent and lateral connections in order to arrive at a simplified view of single-unit neural computation. Even in this simple model though the membrane response of each “AdEx” neuron can be complex spanning chaotic irregular activity, bursting, accelerating, and a decay in rate driven by adaptation (13, 14). Even in the simplest case of regular-firing, with uniform sampling of synaptic weights, a population can exhibit substantial response variability (as shown in Figure 1**a**).

**Fig. 1.**
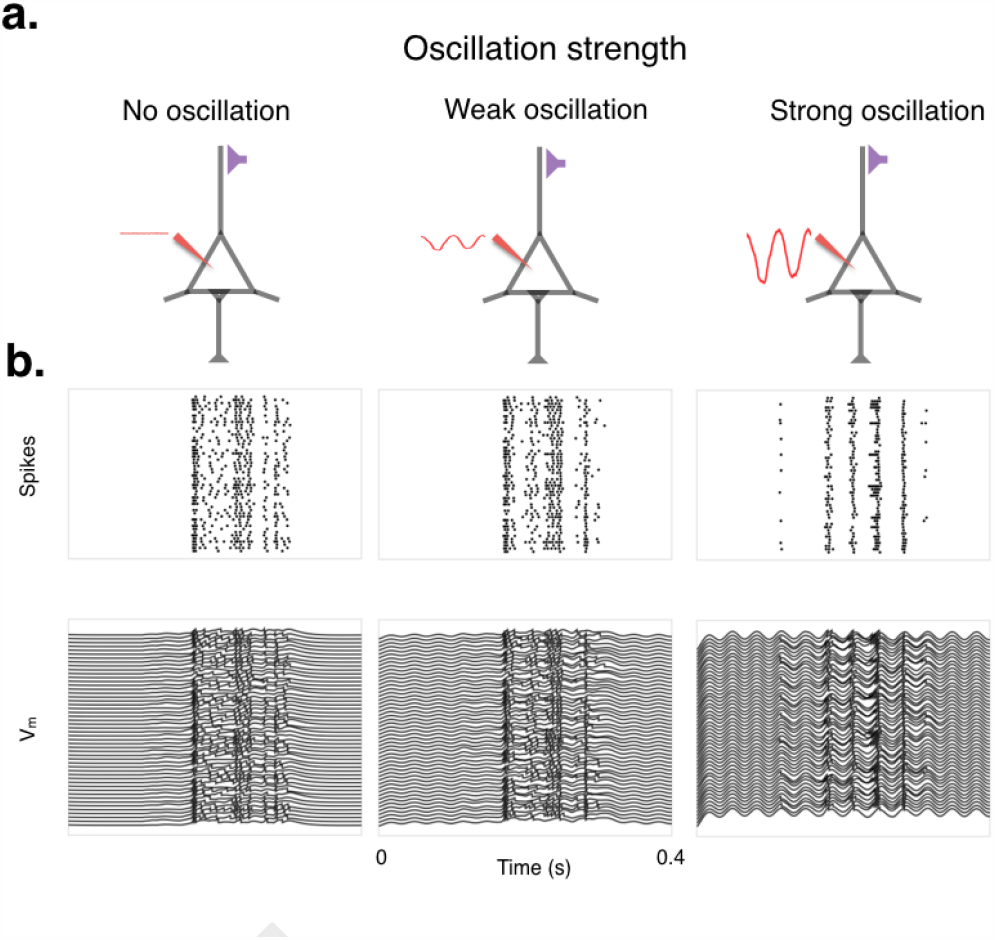
The trade-off between computation and coordination in beta/gamma oscillations. **a**. Illustration of a single model neuron receiving synaptic input (purple synapse), subjected to increasing levels of oscillatory power (red waveform). **b**. A population of *N* independent neurons subjected to three levels of oscillatory power (*A* = 0.0, *A* = 0.1, *A* = 0.6 nS; *f* = 30 Hz). The top panel is a plot of action potentials per neuron and the bottom is membrane voltages of those same neurons.

Brittian *et al* has considered the problem of pathological synchrony in a model of Parkinson’s disease. In line with our illustration, they argued synchrony can tune the complexity of a population’s response. They use this to explain why increased beta (13-30 Hz) synchrony in Parkinson’s patients can lead to a symptom-causing loss of computational capacity (10). While inspirational to our approach, their approach does not offer a way to make quantitative predictions.

In our model a population of independent neurons is entrained by external, global, oscillation in the beta or 20 Hz band. We treat any correlation induced by this entrainment as a perturbation from each neuron’s reference, or aperiodic, computational output—as an error in other words. As the strength of the oscillation grows, then, error increases as neurons are driven to synchronize. By using the term error we do not mean to imply pathology in and of itself. Not all errors are pathological in our view. They are (sometimes) desirable because they lead to synchrony. Instead, we use error to denote a deviation from our reference point. Pathology we define later in terms of degree of error.

Our simulations focus on weak oscillations. We use weak to denote excitatory entrainment that induces little to no new spikes and acts mostly to synchronize spike-times. Though our model is simplified, we aim to mimic the comparatively weak levels of synchrony typically observed in recordings of visual cortex in the beta and gamma bands.

To build a predictive model of the trade-off between synchrony and error in biological terms, those accessible to experimentalists and, perhaps, to the cell itself we examine small temporal moments of membrane voltage. For a small window of time we can separate out the computational (reference) drive from any oscillatory influence, creating what we call a “voltage budget”. That is, how can we “spend” the finite amount of membrane voltage available between the resting potential and action potential initiation. We use these budgets in a predictive way. We estimate the budget at one moment in time to predict how a later immediate cycle of an oscillation will affect spiking.

Our first contribution is the introduction of a voltage budget analysis, which allows us to define a mathematical reasonable, yet biologically observable, criterion to distinguish between healthy and pathological oscillatory synchronization in terms of the membrane potentials alone. We note that even in the healthy range there is a continuous trade-off between the induction of synchrony and the introduction of error. Inspired by observations in our first contribution, we then go on to describe two computer algorithms that can achieve an optimal trade-off between error and synchrony. We finally report that the global oscillator model we study here acts, quite to our surprise, sub-optimally or about two orders of magnitude below the achievable optimal point.

## Results

We model a simple neural network: a population of *N* neurons entrained by a single global oscillator, governed by an amplitude (*A*) and frequency (*f*) of the form *A/*2(1 + sin(2*πf t*) with *A* ≥ 0. When oscillatory amplitude is zero, each neuron is completely independent. As oscillatory amplitude grows, each neuron’s firing is increasingly perturbed by the strength of the oscillation. We illustrate this in Figure 1 where, in the rightmost panel, oscillatory amplitude is relatively high and the population’s firing pattern bears little resemblance to the original population activity as driven by the input (shown in the leftmost panel).

We define computation here in the most basic mathematical sense: an input mapped to an output. We formalize this first in the abstract general form of, *f* : *x* →*y*.. Here *f* is a function, synonymous with computation, and *x* and *y* are any set of inputs and outputs. In practice we implement *f* as an adaptive exponential (AdEx) integrate-and-fire model (15) (see *Methods*), and limit input and output to binary time-series, i.e. action potentials.

### The voltage budget

To understand the boundary between healthy and pathological synchrony, we analyze computation from a neuronal perspective by examining changes in the membrane potential of individual neurons. To simplify this analysis we examine extremely small, nearly instantaneous, windows of time (*w*). For a window of time much less than the membrane time constant (*w << τ*_*m*_) it is reasonable to treat the resting potential (*V*_*r*_) and the firing threshold (*V*_*t*_) as constant values. With these terms fixed, the total amount of voltage becomes fixed and the membrane potential available in each neuron becomes a physically conserved quantity. This means that no energy is allowed in or out, closing the system. To aid our thinking about this conserved system we use economic metaphors. The total amount of voltage available to a neuron in *w* is termed a “budget” that can be “spent” by the neuron. It can be spent to either minimize computational errors—a perturbation of spiking—caused by the global oscillator, or to better align itself with the population (Fig. 2). With this approach the voltage “cost” of increases in oscillatory power can be explicitly analyzed in terms of cellular physiology.

**Fig. 2.**
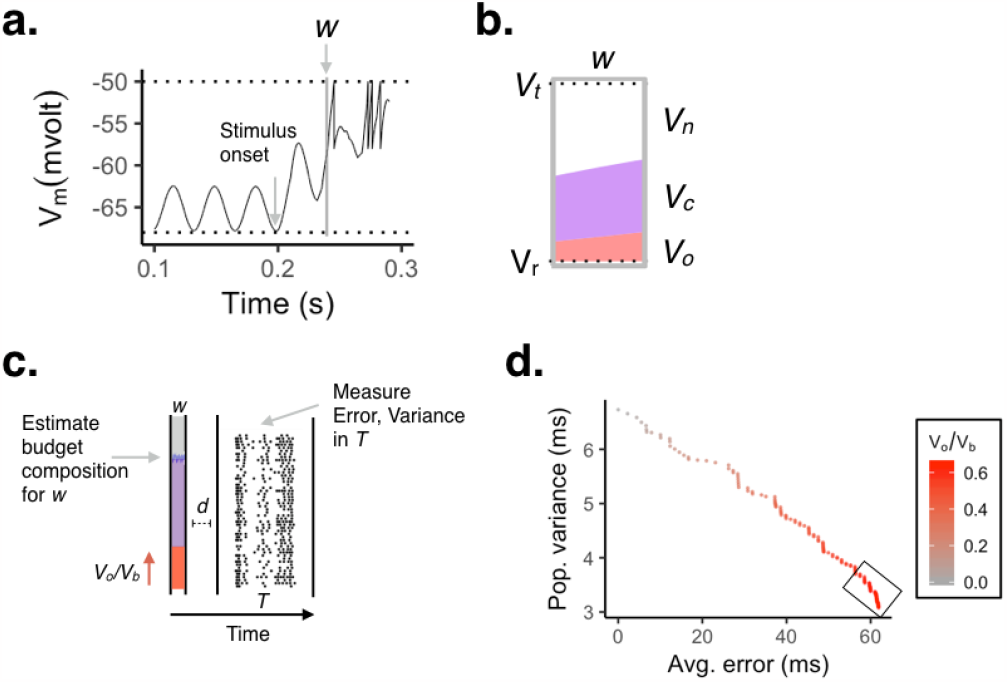
The voltage budget. **a**. Membrane voltage of a single neuron. Horizontal dotted lines indicate the voltage threshold (*V*_*t*_; top line) and the resting potential (*V*_*r*_; bottom line). The vertical gray box depicts the 2 ms budget window (*w*; *A* = 0.1 nA; *f* = 20 Hz). Stimulus onset occurs at 0.2 seconds. **b**. Example of a voltage budget decomposition in time window *w* in panel *a*.. Red represents *V*_*o*_; purple depicts *V*_*c*_; white space is *V*_*n*_. **c**. Diagram of budget prediction. A budget estimate is formed at *t*, then *d* seconds later a single cycle of oscillation begins. Over the period of that cycle’s length *T* we estimate the error and variance of the population spiking. **d**. Plot of average error versus population variance over period *T*. Increases in *V*_*o*_*/V*_*b*_ are denoted in red. Boxed area highlights the error plateau phase. In all plots: *w* = 2 ms; *d* = 2 ms; oscillation frequency *f* = 20 Hz.

We decompose each budget *V*_*b*_ into three terms: 1) the computational voltage *V*_*c*_, 2) the oscillatory voltage *V*_*o*_, and, 3) the open voltage *V*_*n*_ (Fig 1). The open voltage serves the same role as the potential energy term common in basic analyses of the physics of baseball thrown into the air. For example: as the ball rises and is slowed by gravity, kinetic energy is traded for potential energy. Put another way, *V*_*n*_ represents the unused capacity of the system—the difference between *V*_*c*_ + *V*_*o*_ and the threshold potential.

The basic budget relationships are shown in equations 1 and 2. In Eq 2, computation and communication explicitly compete for influence on the eventual spike that happens when the threshold is reached (when *V*_*n*_ → 0).

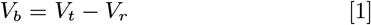

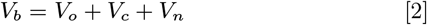

In practice though we study budget terms as ratios, because working in unitless quantities is simpler mathematically and eases empirical comparisons. Specifically, we study the oscillatory power normalized by the total size of each neuron’s budget (*V*_*o*_*/V*_*b*_), and most importantly we study the ratio of oscillation to computation (*V*_*o*_*/V*_*c*_).

### Budget analysis

We imagine each neuron seeks to spend its budget as prudent as possible *for each cycle of oscillation*. We model this by using a voltage budget at time *t* to predict the effect of a later (*t* + *d*) cycle of oscillation on spiking, where *t* + *d* is the time at which we induce a burst of oscillation. We then measure spiking activity for *T* seconds after the oscillation begins, where *T* = 1*/f* (illustrated in Figure 2**c**). Tuning oscillations requires the cell to make temporal predictions, because once a cycle of oscillation begins the dynamics of the network are committed to completing that cycle. Mathematically, oscillations begin as an initial value problem; how the oscillation starts largely determines where it ends.

To demonstrate the broader trend we first examine the effect of a 20 Hz rhythm. Later we show that the key trends for this rhythm is robust to changes to frequency (4-30 Hz), noise (0 - 2.5 mV), and synaptic weights (0.3 -45 nS) (Fig 5), as well as to neuron type (Fig 6).

Each experiment begins with a reference run, where the neurons are initialized with their unique synaptic weights, capacitances, and recovery variables. After 200 milliseconds of settling in time, each neuron is subjected to the stimulus—a pulse of 8 Hz Poisson activity lasting 50 milliseconds. The entire spiking response to this brief stimulus is recorded for later reference. The time of the first spike serves as a locking point *E*. The reference run is followed by the budget run, where we measure the level of computational influence *V* _*c*_. In each neuron, we then calculate *V*_*o*_ for many possible levels of peak oscillatory power *A*, and store these for later reference. We do not imagine that a real biological system would need to explicitly conduct either the reference or the budget phases. This would instead be learned by the system, becoming implicit in the synaptic weights and delay times of the circuit. Finally, in the experimental phase, we sweep over a large range of *A* values corresponding to *V*_*o*_ changes ranging from *<* 0.1 mV to greater than 5 mV. In each of these, a single cycle of oscillation is introduced at *E* +*d* seconds, and spiking behavior is monitored for *T* seconds 2**c**). In all our simulations *d* = 2 ms.

### A model of bursts

We study single cycle “bursts” of oscillation for two reasons. The first reason is empirical: when individual experimental trials of real oscillatory data are examined, oscillations often appear as bursts (16, 17) even though averaging many trials gives what appears to be sustained rhythm (18). As a result of this averaging, most theoretical work on oscillations assume a sustained rhythm though there are exceptions (18–20). Still, it remains unclear what, in general, a single cycle of oscillation can accomplish, which is why we focus on this case here.

The second reason is practical. Bursts, by definition, are accompanied by aperiodic windows of neuronal activity that serve as a natural reference for doing error calculations. By comparing firing during an aperiodic trial to a later periodic trial, we can measure how synchronization perturbs single neuron computations in a simple and direct way.

### The trade-off between error and variance

As *V*_*o*_ rises in proportion to the total budget *V*_*b*_, population variance declines consistent with previous models (21, 22). We also observe that while variance decreases computational errors increase. This trade-off is roughly linear for low power oscillations (Fig2**d**). This early trend implies a nearly one-to-one trade-off between coordination at the population level and errors introduced at the neuronal level. However as relative oscillatory power grows eventually error plateaus at the inflection point of the error-variance curve (boxed region, Fig. 2**d**). Separating the pseudo-linear from the plateau phase is the basis for how we come to separate healthy from pathological synchrony.

### Defining pathological oscillation

The fundamental idea we wish to test: can a pathological oscillation be identified using only computational activity as a reference? To do this we consider the ratio of oscillation to computation, *V*_*o*_*/V*_*c*_. Before getting to our numerical results though it is worth considering *V*_*o*_*/V*_*c*_ on its own. When *V*_*o*_*/V*_*c*_ *<* 1, computation dominates the system in that the majority of the voltage driving each spike reflects the computational dynamics of each neuron. But beyond *V*_*o*_*/V*_*c*_ = 1, the majority of spikes reflect the *modulatory* oscillation. Intuitively then, once a modulator dominates, it is in effect no longer a modulator: the neurons downstream gain information mostly about the oscillatory entrainment “signal” and not the population’s normal computation. This idea leads to our first new definition: oscillations that exceed *V*_*o*_*/V*_*c*_ = 1 are *intrinsically pathological*.

Numerical analysis of *V*_*o*_*/V*_*c*_ confirms this prediction: at *V*_*o*_*/V*_*c*_ = 1.0, the average error and variance of the population plateau and are incrementally—asymptotically—approaching their maximum values (for error see Figure 4**a** and **c**). At the individual level, error for all neurons plateaus by *V*_*o*_*/V*_*c*_ = 1.0, however the steepness and curvature of this rise is neuron dependent (Figure 4**b**). These cell-level differences suggest that global oscillation is not optimal. That is, better trade-offs may be possible if the oscillation is tuned to each neuron’s particular computational curve; an idea we return to below.

### Defining healthy oscillation

In a later section we prove that the optimal trade-off between error and variance is linear. This proof leads to our second criterion: on normative grounds we suggest that healthy oscillations have *V*_*o*_*/V*_*c*_ *<* 0.5. This point demarcates the average transition from pseudo-linear to plateau (Figure 3**a** and **b**). At *V*_*o*_*/V*_*c*_ = 0.5, this point also separates the transition in variance from linear to its (even sharper) plateau phase, after which increasing oscillatory power has an increasingly marginal effect on error or variance, as those are close to maximal and minimal respectively (3**c**).

**Fig. 3.**
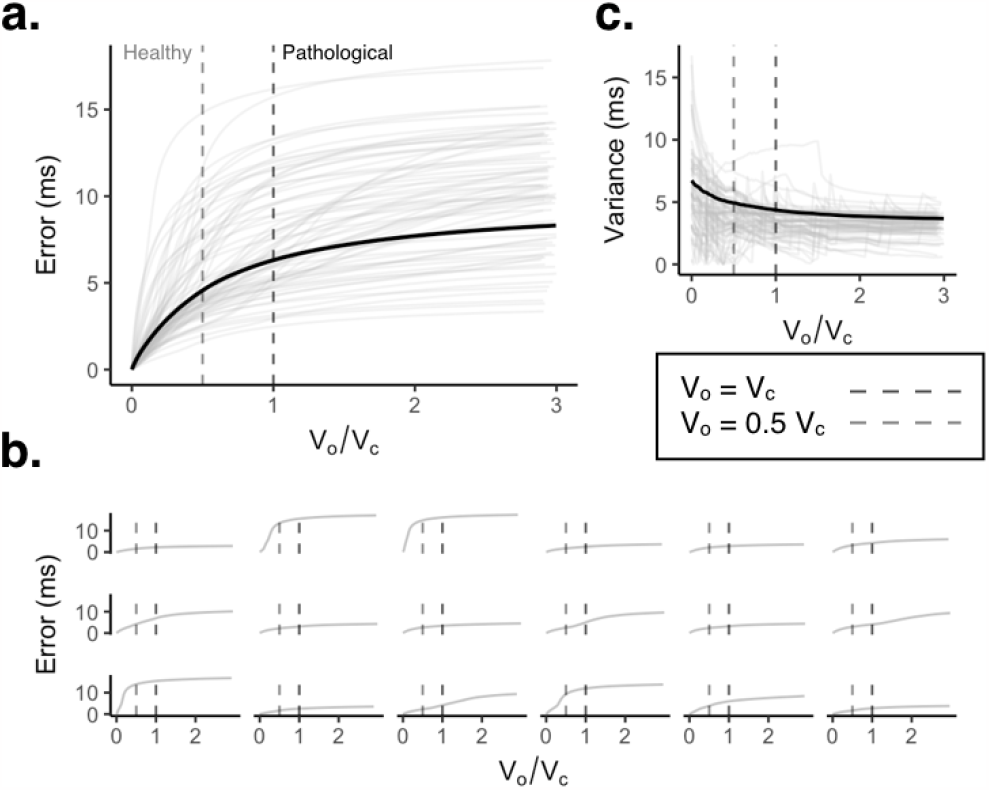
Voltage budget spent on beta/gamma oscillations increases error and reduces variance. **a**. Plot of error predicted by changes to *V*_*o*_*/V*_*c*_. Gray lines indicate individual neurons. Black is the population average. Dashed lines suggest the end points for the pathological (dark) and healthy (light) inequalities. **b**. Randomly chosen individual examples taken from *a*. **c**. Variance predicted by the ratio *V*_*o*_*/V*_*c*_. Gray lines indicate individual neurons. Black represents the population average.

### Variance and

*V*_*o*_. Variance and oscillatory budget (*V*_*o*_) have a more complex relationship than error and oscillation. In individual neurons we see that as oscillatory budget increases variance exists as two phenomena, which act in opposition. For sufficiently small increases in oscillatory budget, what we term a “weak” oscillation, synchrony declines along a logistic path (see light orange traces in Figure 4**a** and **e**). However once the oscillatory budget reaches a “strong” critical value–that is specific to each neuron–variance discontinuously increases, denoted by the dark orange in Figure 4**a** and **e**. These sudden jumps happen every time a new, extra action potential is generated. New action potentials tend to be at the extremes of an oscillatory cycle, as illustrated in Figure 4**f**. In contrast to variance, however, individual neuron’s errors change smoothly (Figure 4**e**).

**Fig. 4.**
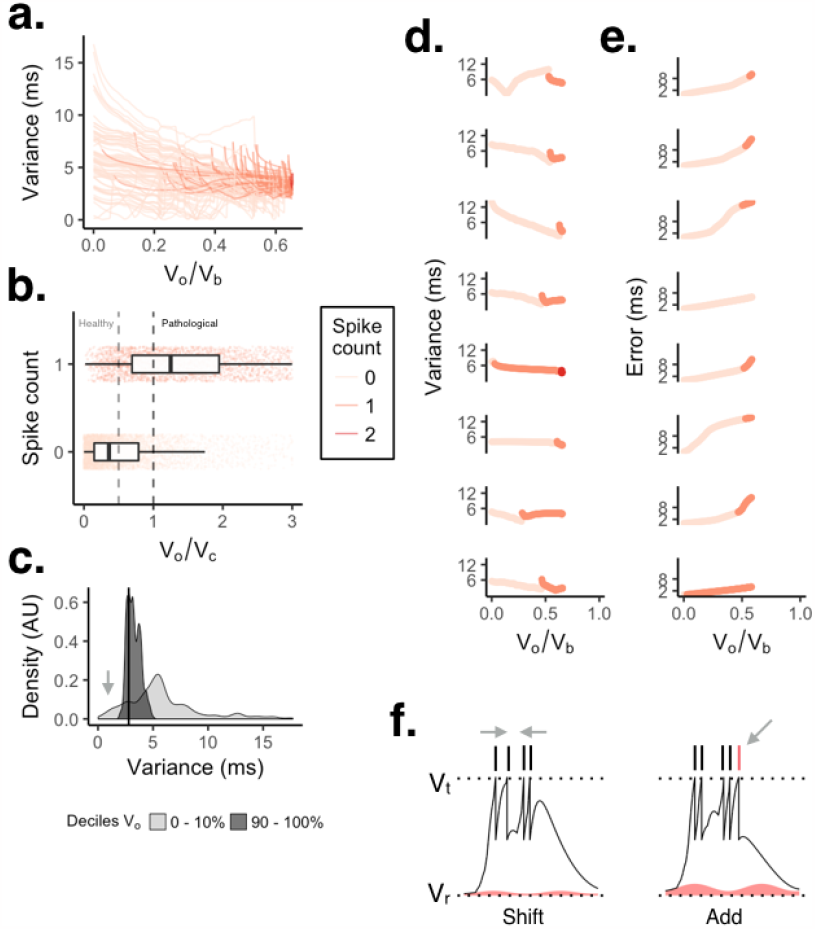
Oscillations both decrease and increase variance depeding on if they are “weak” or “strong”. **b**. A plot of variance and *V*_*o*_, where traces from individual neurons are colored by the number of new spikes introduced by *V*_*o*_. Light orange indicates a weak oscillation that introduced zero new spikes; dark orange is when a strong oscillation introduced one spike; red is two spikes. **b**. Spike count as a function of *V*_*o*_*/V*_*c*_. The left side of the light dashed line denotes the healthy ratio of *V*_*o*_ to *V*_*c*_ while the dark line denotes the start of the pathological region. **c**. Histogram density plots for the top and bottom variance deciles. The black line indicates the minimum population variance. The arrow indicates neurons that show a lower total variance for weaker, rather than stronger, oscillations. **d**. Randomly chosen examples of variance for individual neurons. **e**. Examples of error for individual neurons. **f**. Illustration of the opposing possible effects an increase in *V*_*o*_ can have on variance. Left panel depicts an oscillation acting to shift spike times closer together. Right panel depicts a stronger oscillation that adds an additional spike, shown in red.

On average, increases in oscillatory budget increase synchrony. Counter-intuitively, however, very weak oscillations can generate as much synchrony as strong oscillations. In fact, the quantiles analysis in Figure 4**c** suggests that weak oscillations, when targeted at a select subset of neurons, can have lower overall variance, while showing much greater capacity for error/variance trade-off. This targeted phenomenon is highly consistent with recordings of gamma oscillation in visual cortex, where only a few neurons appear to be coupled to the oscillatory phase (**?**).

### Testing robustness to frequency

Frequency has little effect on error (Fig. 5**a**) but profoundly affects the synchrony (Fig. 5**b**). Oscillations slower than 20 Hz show similar trends outlined in Figures 2-5 (yellow arrow, Fig. 5**b**), while oscillations faster than 20 Hz generate less total synchrony and plateau faster (green arrow, Fig. 5**a**).

**Fig. 5.**
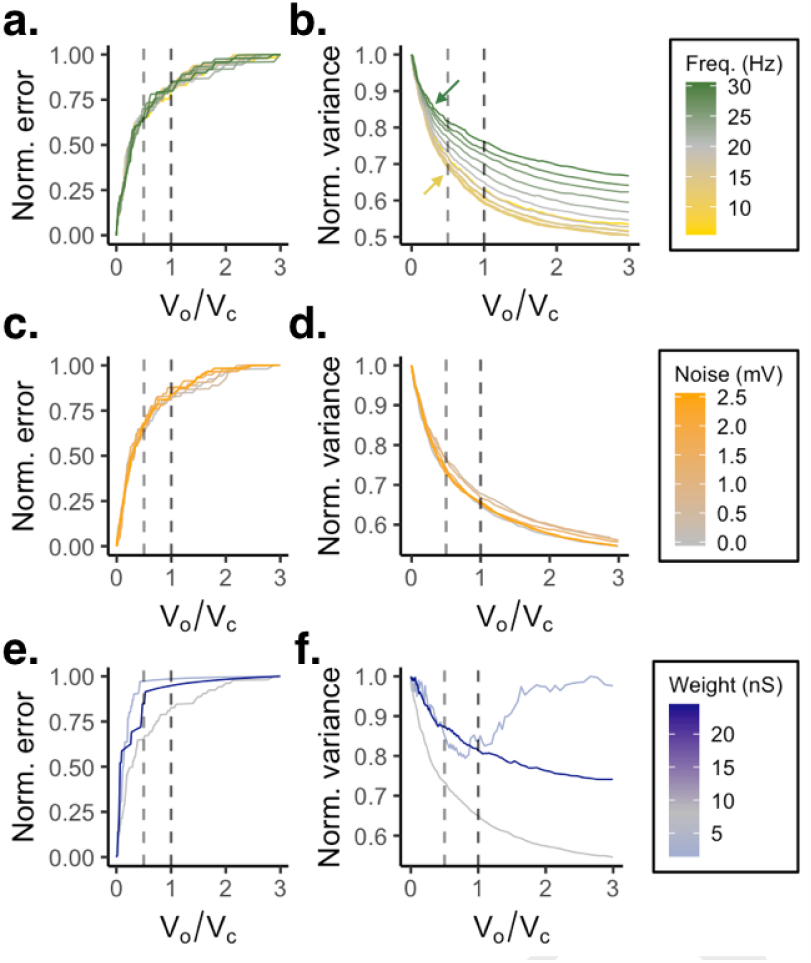
Testing oscillatory budget robustness to frequency, noise, and synaptic weight. The gray colored curve on each plot represents the “standard” value otherwise used throughout. Light and dark dashed lines represent healthy and pathological *V*_*o*_*/V*_*c*_ ratios, respectively. **a**. Simulated effect of oscillation frequency (4-30 Hz) on average error, normalized against each neuron’s maximum error. **b**. Population variance decreases more quickly for stronger oscillations when those oscillations are lower frequency. **c**. Injection of noise into membrane voltage between 0 and 2.5 mV shows that noise has little effect on either error or, **d**., variance as a function of oscillatory power. Oscillation frequency was 20 Hz (gray curve). **e**. Simulating changes in the synaptic weight range and its impact on average error. Plots are colored by the average weight. Individual ranges for the standard weight value in gray were 1.5-15 nS. Light blue illustrates weak synapses in the 0.3-3 nS range, while dark blue represents strong synapses sampled from 4.5-45 nS. Individual synaptic weights were sampled independently and uniformly for each condition. Oscillation frequency was 20 Hz (gray curve). Synaptic weights have relatively weaker effects on population error, but, **f**., show stronger effects on population variance.

### Influence of membrane noise and synaptic weight

Biologically-relevant levels of noise have little effect on normalized variance or error (Fig. 5**c**-**d**). The range of synaptic weights, in combination with the input firing rate, modestly alter the normalized error profile (Fig. 5**e**) but can strongly affect variance (Fig. 5**f**). As shown in Fig. 5**f**, when operating in the healthy range, where *V*_*o*_*/V*_*c*_ ≤0.5, all examined synaptic weights show a consistent monotonic decrease in synchrony as *V*_*o*_ rises. However in the pathological range *V*_*o*_*/V*_*c*_ ≥ 1, weaker synapses and high *V*_*o*_ generate a strong increase in variance—a change in variance 5-fold larger than any other observed in our simulations. The strength of this effect might tempt us to speculate that weak synapses are particularly prone to generating pathological symptoms. When placed under what might otherwise be considered a low level of power and weak level of entrainment, neurons with weaker synapses may generate unusually asynchronous responses.

### Influence of cell-type

So far we have focused on regular firing neurons. Now we consider a heterogeneous population of neurons displaying a wide range of firing modes. Our neuron dynamics are governed by the AdEx model, which is known to fit a wide range of real firing properties including fast firing, adaption, accelerating activity, bursting, and chaotic irregular firing (14). However, rather than simulating preset classes of neurons, we opted instead to smoothly sample a large range of the neural space, generating a diverse pool of firing behaviors.

The space is illustrated by a few randomly selected examples in Figure 6**a**.

On average, the same budget ratios that predict healthy synchrony in regular firing neurons hold true in simulations of heterogeneous neuronal populations Fig. 6**b**-**c**). All models show the same one-to-one trade-off between variance and error below *V*_*o*_*/V*_*c*_ = 0.5 and an error plateau by *V*_*o*_*/V*_*c*_ = 1.0.

**Fig. 6.**
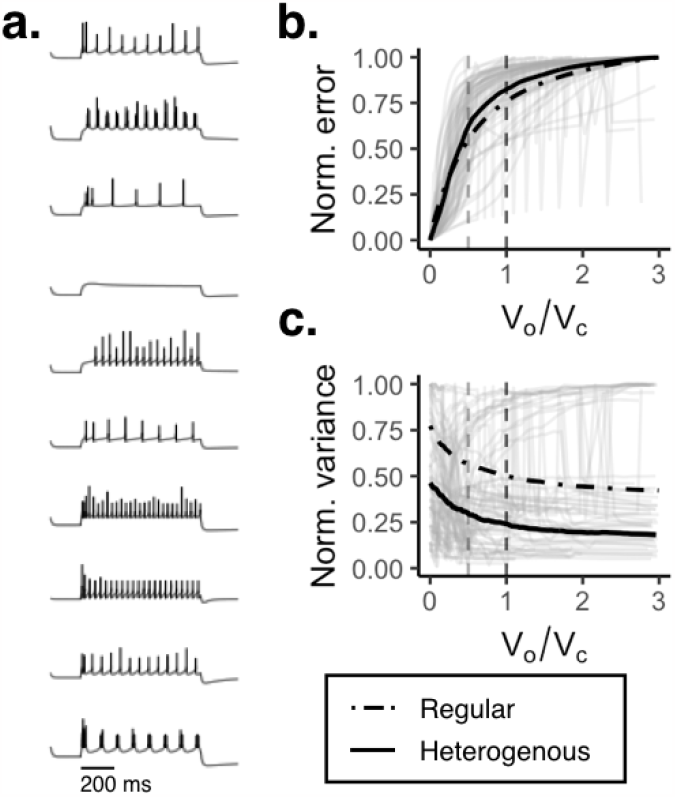
Influence of cell type on voltage budget analysis. **a**. Examples of heterogeneous firing modes, driven by a 0.8 second, 0.25 nA square-wave pulse. **b**. Average (black) and individual errors (gray) for *N* = 250 heterogeneous neurons. Average error from a regular firing network is redrawn from Figure 3*a* (dot-dash). The left side of the light vertical line denotes the healthy oscillation ratio ((*V*_*o*_/*V*_*c*_)=0.5) while the dark light denotes the start of the pathological region ((*V*_*o*_/*V*_*c*_)=1.0). **c**. Variance as a function of the *V*_*o*_*/V*_*c*_ ratio. Black is the population average. Grey traces represent individual neurons, and the dot-dashed line represents variance from the regular population.

### Optimal synchronization

To measure population synchrony and computational error we have used two related metrics. The first is the mean absolute error (*E*), which measures the average computational error. The second is the mean absolute deviation (*D*), which measures the variance of each neuron spiking relative to the population average. Exploring the mathematical connections between these metrics offers insight into the structure of the error/synchrony trade-off, and allows us to prove two optimal algorithms for spike-time synchronization.

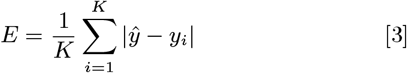

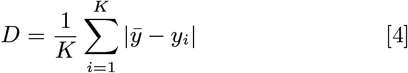

Here *y* are spike times from individual neurons, reflecting an example when oscillatory amplitude *A* is greater than zero. *ŷ* is the set of reference times acquired without oscillations, *i*.*e*., when *A* = 0. We denote examples from *y* as *y*_*i*_, and 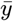 is the average of *y*.

What’s the best algorithm to shift spike times? To formalize an answer, assume we wish to change spiking variance *D* by some amount *ϵ* ∈ R^1^. When *ϵ* is 0 there is no change in synchrony and so by definition *y* = *ŷ* and *E* = 0. As we increase *ϵ* the question becomes how should we distribute that perturbation, or error, among the K spikes in the spiking population *y*? That is, how do we set each neuron’s spiking error, *ϵ*_*i*_, for the series *y* = (*ŷ*_1_ + *E*_1_, *ŷ*_2_ + *E*_2_) … (*ŷ*_*K*_ + *E*_*K*_)?

A naive approach, similar in character to a global oscillator, is to spread the error uniformly among the entire spiking population, *y*. Formally, if we decompose total error into *i* equal error partitions, then we have a uniform error distribution case where *ϵ* = |*ϵ*_1_| + |*ϵ*_2_| + … + |*ϵ*_*K*_|. If we wish to use error to minimize variance we must set the sign of each perturbation *ϵ*_*i*_ to oppose the sign of *y*_*i*_; if *y*_*i*_ is negative, *ϵ*_*i*_ is positive, and vice versa (for example see Figure 7**a**, *bottom panel*). When implemented over a range of variances, this uniform approach gives rise to the blue error-variance curve in Figure 7**b**. The question then becomes is this uniform strategy the optimal algorithm to balance the trade-off between computational error and population variance? That is, is there a smaller value of *D* for a given level of error, *E*?

**Fig. 7.**
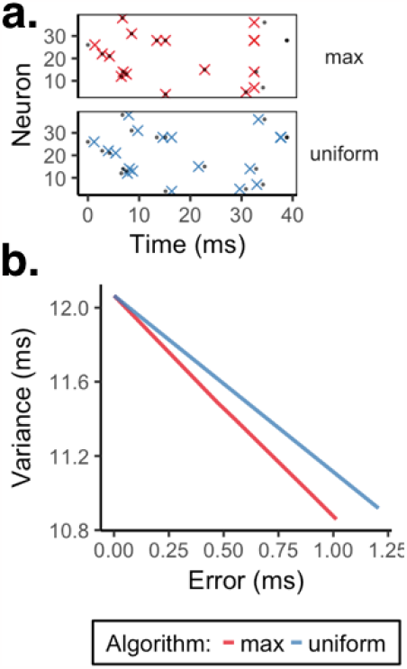
A comparison of two algorithmic strategies for inducing synchrony. **a**. In these examples, two populations of synchronized neurons have their original spikes (•) shifted to new positions (X) based on two different algorithms: either by iteratively shifting the spikes furthest from the mean (top, red) or by shifting each spiking uniformly (bottom, blue). **b**. Error-variance trade-off curves for the strategies illustrated in *a*. The smaller the error for a given level of variance, the more optimal the algorithm is. Note that the *max*(|*y*|) algorithm (red) optimally minimizes the trade-off between computational error and population variance.

To explore optimality, we introduce a single degree of freedom. We hold all errors equal as before, except for two neurons *m* and *n*. This lets us ask the question: by introducing a single degree of freedom can we generate more synchrony than the simple uniform error distribution strategy? If we can do so, we know that the uniform strategy is *not* optimal. To simplify the analysis, first we center all *y* and *ŷ*, by subtracting 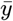 from all, and remove the normalization term 1*/K*, leading to equations 5 and 6.

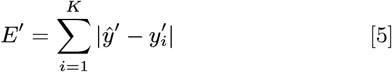

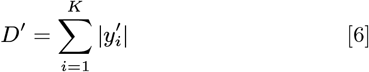

An instructive, but extreme, use of a single degree of freedom is to assign all the values from one free perturbation to the other. That is we set *ϵ*_*n*_ to 0, and *ϵ*_*m*_ to 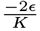. From here it becomes clear that if we apply *ϵ*_*m*_ to max(|*y*|) and *ϵ*_*n*_ to min(|*y*|) this will produce the largest possible decrease in adjusted variance, *D*^*I*^, and represents a max(|*y*|) min(|*y*|) decrease in *D*^*I*^ compared to the uniform approach.

If we free ourselves from the contrived example of uniformly distributing *ϵ*, we can see now that the optimal approach is to apply *ϵ* to max(|*y*|); max(|*y*|) → max(|*y*|) ± *ϵ*. This means that for any given arrangement of spike times, the optimal trade-off between computation and population synchronization introduced by an oscillation is to shift the spike farthest from the mean. We implement this as an incremental algorithm, which can change the variance by *ϵ* by intuitively ranking *y*, shifting the spike max(*y*) by some very small amount in time *δ* until a running sum of all these tiny perturbations is equal to *ϵ*. In theory, the smaller the *δ* the better the algorithm approximates the true optimal solution. In practice, a *δ* below 0.00001 seconds is sufficient. See Figure 7**a** (*top panel*) for an example of a synchronized spike train and **b** for a depiction of the optimal error-variance curve.

### Global oscillation is very suboptimal

Having derived two optimal algorithms to trade error for variance, we compared these to our oscillator model (Figure 8). We first ran the oscillator model–in regular firing mode–over a range of weak oscillations (*A* = (0, 0.02*e* − 9), *N*_*A*_ = 150). We focused on weak oscillations because they generate the lowest overall variance per error; a best case scenario. For each power *A* we recorded spike-times, their error and variance. We then ran our optimal max deviance and uniform algorithms on these same spikes times, producing optimal error values. The error curves from the oscillator, max deviance, and uniform are the shown Figure 8).

**Fig. 8.**
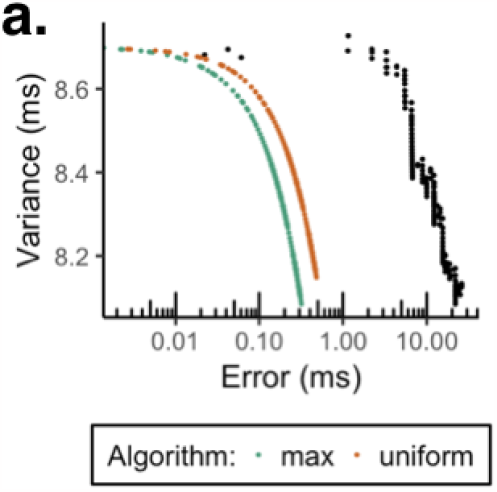
A comparison of optimal error-variance curves found using our max deviance (shown in teal), or the uniform (or max min, orange) algorithms compared to that introduced by a global oscillator (black). Oscillation frequency was 8 Hz. See Figure 7 for more detail about these two algorithms.

The global oscillator produces errors that are 1-2 orders of magnitude larger than either of the optimal value. Other runs, with different stimuli, synaptic weights, and cell firing types, have qualitatively identical results (not shown). So while our model of a global oscillator does synchronize firing, this leads to highly suboptimal increases in error. If that is, the proper error reference is found population without oscillation

## Discussion

We build a simple model of external oscillatory entrainment. We mimic the real biological case where one “pacemaker” population coordinates another, aperiodic, population, such as in the case of top-down oscillatory influence (23–26). Our model is the simplest case we could devise that allows for the precise, biologically testable, predictions of oscillatory over-entrainment.

### Limits of the model

We studied a toy model designed as a best-case scenario for understanding the trade-off between computation and communication. Our purpose was to create an initial (the first, as far as we are aware) quantitative model of healthy versus pathological oscillations, and so begin by using the simplest model that embodies the problem: uncoupled neurons subjected to a global oscillator. Real biological systems, and more complete simulations, feature extensive yet sparse connections between neurons. These connections naturally create dependencies between the activity of one neuron and the others in its population—a basic phenomenon we do not capture here. Theoretical analysis of neural coding, and decades of empirical research, however, suggest that, despite this extensive connectivity, real neurons act with a high degree of independence, which is optimal for computational efficiency. The biological implementation of this independence relies on a combination of independent dendritic computation and the precise arrangement of excitatory and inhibitory circuits. So, despite the simplicity of our model, it may act as a reasonable approximation of real complex networks which maintain a high degree of independence between neurons.

Oscillations in a network can also arise by self-organization, where rhythmicity is driven by interactions *within* the entrained population. Our model is not well-suited to this case; previous theoretical work suggests that, even in idealized cases, there is a minimum level of voltage budget needed to initiate and sustain an intrinsic oscillation (27). That is, in self-organized systems the order parameter *can’t* be expected to smoothly vary in the biologically relevant 1-5 mV range, which is a requirement for our analysis to hold. Understanding the interaction between computation and coordination within a self-organized population remains an open question.

### Real synchrony is often weak

It is not clear how oscillatory entrainment can be weak, yet also be an important general feature of nervous system function, as is frequently supposed (2, 3). That is, to observe oscillations in real local field potentials often requires little more than placing an electrode in the appropriate region, as field potential oscillations are relatively ubiquitous (24, 28). Observing the same oscillation in the spiking behavior of neurons, however, often requires recording from many—even hundreds—of neurons, especially in cortical areas (29–31). In these recordings about half the neurons show no preference for an oscillation’s phase (32). Those that are entrained are often weakly entrained, synchronized by at most few percent.

Our model suggests that the most effective oscillations are precisely those that are both weak and sparse. When the oscillatory power remains below the that of all other “computational” inputs, the system can exchange errors in a single neu-ron’s computation for group-level synchrony, measured in the voltage budget analysis as the quantitative ratio *V*_*o*_*/V*_*c*_ ≤0.5.

This ratio’s predictions are relatively invariant to oscillation frequency, noise in the membrane potential, and variations in synaptic weight; this ratio also predicts the firing properties of a large range of heterogeneous cell-types. Further, strong oscillations offer only marginal improvements in synchrony: once an oscillation grows too strong it induces new action potentials in the population. These extra action potentials tend to be at the trailing end of the neuron’s response to input, increasing variance rather than gathering spikes together. Finally, our new strategy for provably-optimal coordination targets a small fraction of the population. Targeting all neurons for coordination has a larger error cost than targeting only a few of the more extreme action potentials in a given cycle.

### Oscillations as epiphenomenon

Oscillations could be a side effect, or epiphenomenon, of neural physiology. Mathematical and experimental analysis of both simple (30, 31) and complex biological (13) structures suggests a relatively large portion of the neural parameter-space generates oscillations. As a result, oscillations may be only a nuisance. An artifact of other biological factors. An epiphenomenon.

On the other hand, oscillations may have arisen early on during nervous system evolution, initially as an artifact. However, over time these oscillations were co-opted and put to, perhaps several distinct, uses. After more than 80 years of study, separating these two possibilities remains an open problem.

We offer a new approach to the oscillations-asepiphenomenon debate. By deriving *a priori* quantitative bounds between the healthy and pathological ranges of oscillation, and in defining an optimal algorithmic approach to synchrony, we suggest that these normative constraints can help in finally separating functional oscillations from physiological epiphenomena. To see how, recall that our analysis suggests that oscillatory input—and all the other neuronal inputs (collected into the “computational” term in our model)—into the neuron exist in equilibrium. At one end of this equilibrium are neurons whose action potentials are independent. At the other end are neurons who are completely synchronized, and therefore redundant. Oscillations that are *just* an artifact would be expected to explore both extremes. On the other hand, oscillations that track strict normative bounds must be functional, rather than epiphenomenal.

## Methods

### The network

Our model was a network of *N* = 250 Adaptive Exponential (AdEx) neurons (15) sharing a common oscillatory drive, 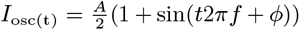 (1 + sin(*t*2*πf* + *ϕ*)); where *f* is the frequency of the oscillation, and *ϕ* denotes the phase offset. Each neuron was driven by an identical Poisson process. Variability in computational output depended solely on the variations in synaptic weight, and ‘slow” membrane recovery parameters *a* and *τ*_*w*_, all of which were independently sampled from a uniform distribution (15) (Eq. 9).

All synapses were excitatory, and governed by a single exponential decay term with time constant *τ*_*e*_. Membrane noise currents were driven by a Ornstein–Uhlenbeck process, with a 5 ms time constant.

Formally the synaptic input, *I*_*x*_, consisted of a spike train *x*, Poisson-distributed (*λ* = 8), a synaptic weight *w*_*x*_, and passive conductance *g*_*x*_.

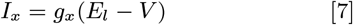

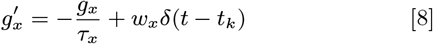

Membrane dynamics were governed by an AdEx model (Eq. 9).

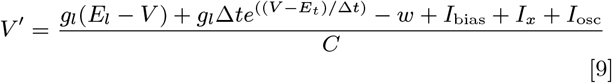

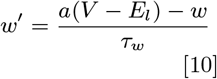

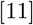

Here the leak conductance *g*_*l*_, the capacitance *C*, and the leak potential *E*_*l*_ control the passive properties of each neuron. While action potential initiation is driven by the exponential Δ*te*^(.)^ (Eq 9). Following an action potential, the membrane has both a “fast” (instantaneous) step, where *V* and *w* are reset (Eq. 12), and a “slow” (passive) response governed by dynamics of *w* itself. Here *a* and *τ*_*w*_ define the rate of change of *w* (outside of “fast” events).

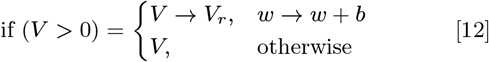

### Neuron-type

Initially we modeled regular firing neurons, whose parameters were published in (14). To generate the heterogeneous population in Figure 6, we sampled uniformly from within a large range of AdEx membrane parameters, whose values are found in Table 1. These were in turn drawn from several neuron types previously described by (14).

**Table 1.** Model parameters.

| Symbol | Range (unit) | Description |
| --- | --- | --- |
| $A$ | 0 - 0.15 (namp) | Peak current (order parameter) |
| $\phi$ | 0 | Oscillation phase |
| $f$ | 0 - 30 (Hz) | Oscillation frequency |
| $r_s$ | 6 (Hz) | Stimulus firing rate |
| $C$ | 100-200 (pfarad) | Membrane capacitance |
| $w_{\text{in}}$ | 1.5 - 15 (nsiemens) | Synaptic weight |
| $\tau_e$ | 5 (msecond) | Excitatory synaptic time constant |
| $V_e$ | 0 (mvolt) | Excitatory synaptic reversal potential |
| $I_{\text{bias}}$ | 10 (namps) | Bias current |
| $\sigma$ | 0 - 5 (namps) | Injected membrane noise |
| $g_l$ | 10 - 18 (nsiemens) | Leak conductance |
| $V_l$ | -50 - -70 (mvolt) | Leak reversal potential |
| $a$ | -11 - 4 (nsiemens) | Slow recovery term |
| $\tau_w$ | 30 - 300 (msecond) | Slow time constant |
| $b$ | 0 - -12 (psiemens) | Instantaneous recovery term |
| $V_{\text{rheo}}$ | -46 - -58 (mvolt) | Instantaneous recovery voltage |
| $\Delta t$ | 2.0 (msecond) | Recovery voltage rate term |

That is, rather than simulate a pre-set selection of specific neurons, during heterogeneous experiments each neuron’s parameters were sampled from a uniform distribution bounded by the extreme values found in (14). For example, the smallest capacitance *C* found by Naud *et al* was 100 picofarads. The largest was 200. This means in our heterogeneous model each *i*th neuron’s capacitance *C*_*i*_ was independently sampled as *C*_*i*_ ∼ *U* (100, 200). Continuing this pattern, the leak conductance *g*_*l*_ and voltage *V*_*l*_ were sampled from (10, 18) nS and (-59, -70) mV respectively. The slow recovery dynamics controlled by *a* and *τ*_*w*_ were sampled from (-11, 4) nS and (30, 300) ms. The instantaneous after-spike reset parameters were *b*: (0.0, 120) picoamps and *V*_rheo_: (-46, 58) mV.

All model parameters are summarized in Table 1.

### Metrics

Computational error and synchrony were estimated with a related set of metrics—the mean absolute error *E* (Eq. 3) and mean absolute deviance (*D*, Eq. 4). In its mathematical form the mean absolute error assumes the reference *ŷ* and observed variables *y* to be the same size. In practice here, this constraint could not be satisfied; as *A* increases new action potentials are inevitable. To accommodate potentially uneven lengths, we truncated the longer of the two to match the shorter. For example, if the target series had length *K*, and the observed sequence had length *L*, when *K > L, K* − *L* elements were removed starting with the largest value. When *L > K, L* − *K* of the largest were eliminated.

## ACKNOWLEDGMENTS

B.V is supported by a Sloan Research Fellowship, the Whitehall Foundation (2017-12-73), and the National Science Foundation (1736028). EJP wishes to thank Rachel L. Storer for invaluable discussions during development of the budget analysis approach.

## Notes

The authors have no conflicts of interest to declare.

https://github.com/parenthetical-e/voltagebudget

### Competing Interest Statement

The authors have declared no competing interest.

### Summary of Updates

Minor changes to framing to align with its companion paper found at, https://www.biorxiv.org/content/10.1101/2021.04.24.441276v2

